# Gonadal sex and sex chromosomes each contribute to sexually dimorphic gene expression in threespine stickleback

**DOI:** 10.64898/2026.05.12.724688

**Authors:** Matthew J Treaster, Michael A White

**Affiliations:** Department of Genetics, University of Georgia, Athens, GA 30602

## Abstract

Many taxa have evolved heteromorphic sex chromosomes like the XY system found in mammals. In additional to the sex determination gene which directs development of the gonad into an ovary or testis, sex chromosomes can have drastically different gene content, leading to substantial genetic differences between genetic males and females beyond their gonad identity. Studying the effects of these genetic differences is challenging, as the sex chromosomes and sex determination gene are inherited together, so the effects of genetic differences between the X and Y cannot be easily isolated from the hormonal differences produced by the ovary and testis. The threespine stickleback fish has a heteromorphic XY sex chromosome system and a wide range of well documented sex differences in morphology and behaviors, including complex mating behaviors and male-only parental care. Through genetic manipulation of *amhy*, the newly identified sex determination gene in threespine stickleback, we are able to generate gonadal males and females with either the XX or XY sex chromosome complement and analyze the separate effects of gonadal sex and sex chromosome complement on sexually dimorphic gene expression. We find that sex chromosomes have a larger effect on gene expression than gonadal sex in somatic tissues, while gonadal sex has a larger effect on expression in the gonads. We also find that the X and Y chromosomes are enriched for genes which show differential expression between females and males. Our findings demonstrate the significant biological impact of sex chromosomes outside of primary sex determination and showcase the utility of the threespine stickleback for studying the genetic basis of sex differences.

## Introduction

Sex chromosomes have evolved countless times across eukaryotes [1–4]. Sex chromosomes typically begin as a pair of autosomes which recombine freely and are otherwise identical until one of the autosomes acquires a SD gene [1,4,5]. In some instances, an SD gene can exist stably without further changes in the surrounding region [4]. In many cases, recombination is suppressed around the sex determination gene, causing the chromosomes to differentiate [1,4,5]. This non-recombining region may later expand in a stepwise fashion, forming multiple strata of different ages [1,4,5]. The cause of recombination suppression is not clear, but the most widely accepted model proposes that recombination suppression is selected for to link the SD gene with nearby sexually antagonistic loci that are beneficial to one sex but detrimental to the other [2,4,5]. Although this model is challenging to test, and empirical evidence supporting it is sparce [6,7], the widespread prevalence of highly diverged, heteromorphic sex chromosomes implies that this phenomenon provides some evolutionary advantage.

There are some known traits with that are influenced by sex chromosomes outside of their contribution to sex determination. Sexually antagonistic color morphs are associated with the Y chromosomes in guppy [8,9] and W chromosomes in two clades of cichlids [10,11]. In *Drosophila melanogaster*, Y chromosome variation influences locomotor activity [12] which has been shown to experience sexually antagonistic selection [13]. Identifying traits influenced by sex-linked loci is challenging in many species as the effects of the SD gene and hormones produced by the gonad cannot be disentangled from the effects of other genes on the sex chromosome. Such observations typically rely upon standing variation between sex chromosomes across populations, use of exogenous hormones to induce expression of sex-limited traits, or rare, naturally occurring sex reversed individuals. In mouse, researchers have manipulated the SD gene, *Sry*, to generate male (having testes) and female (having ovaries) of either sex chromosome complement (XX or XY) [14], allowing the separate effects of sex and sex chromosomes to be quantified. This “Four Core Genotypes” mouse model has illuminated the effects of sex chromosomes on an astounding range of traits (reviewed in [15,16]). Unfortunately, no similar systems exist outside of mammals, precluding comparative research in this field.

The threespine stickleback fish (*Gasterosteus aculeatus*) is a powerful emerging model with a history of research in both sex chromosomes [17–21] and sex differences [22–43]. We recently identified a Y-linked copy of anti-Mullerian hormone (*amhy*) as the SD gene in the threespine stickleback [44]. Using CRISPR/cas9 and a transgene containing *amhy*, we generated sex reversed XX males and XY females which had fully developed gonads and viable gametes. This finding presented an exciting opportunity to study the separate contributions of gonadal sex and sex chromosomes to sexual dimorphism in a novel system. In this study we analyze the transcriptomes of brain, gill, gonad, and liver in juvenile wildtype and sex reversed stickleback. We find that sex chromosomes have a larger effect on gene expression than sex in somatic tissues, while sex has a larger effect than sex chromosomes in the gonad. Both the X and Y chromosome are enriched for genes that show sex differential expression in somatic tissues. These findings show that sex chromosomes can have significant biological impacts outside of their role in regulating primary sex determination, demonstrating the utility of the Four Core Genotypes stickleback for studying the genetic basis of sex differences.

## Results

### Study Design

We analyzed gene expression in gonadal males and females with either XX or XY sex chromosomes. Sex reversed XY Females (XY-F) carry a previously described frameshift *amhy*-KO allele generated with CRISPR/Cas9 [44]. Sex reversed XX Males (XX-M) contain a transgene with *amhy* under control of its putative native regulatory elements [44]. We sacrificed fish at 200 days post hatching. Fish were between 31 and 38 mm in standard length (Table 3.1), and there was no significant difference in length between genotypes (ANOVA, p = 0.30). We collected and sequenced transcriptomes from brain, gill, gonad, and liver tissue from six replicates for each genotype.

**Table 3.1.** Standard length of fish at the time of tissue collection.

| Sample | Sex chromosome | Gonadal Sex | Standard length (mm) |
| --- | --- | --- | --- |
| XX-F-1 | XX | Female | 31.5 |
| XX-F-2 | XX | Female | 35 |
| XX-F-3 | XX | Female | 34.5 |
| XX-F-4 | XX | Female | 35.5 |
| XX-F-5 | XX | Female | 37 |
| XX-F-6 | XX | Female | 33.5 |
| XX-M-1 | XX | Male | 36 |
| XX-M-2 | XX | Male | 35.5 |
| XX-M-3 | XX | Male | 36.5 |
| XX-M-4 | XX | Male | 34 |
| XX-M-5 | XX | Male | 36.5 |
| XX-M-6 | XX | Male | 36.5 |
| XY-F-1 | XY | Female | 35.5 |
| XY-F-2 | XY | Female | 33 |
| XY-F-3 | XY | Female | 36.5 |
| XY-F-4 | XY | Female | 34.5 |
| XY-F-5 | XY | Female | 32.5 |
| XY-F-6 | XY | Female | 31.5 |
| XY-M-1 | XY | Male | 37 |
| XY-M-2 | XY | Male | 38 |
| XY-M-3 | XY | Male | 36 |
| XY-M-4 | XY | Male | 34 |
| XY-M-5 | XY | Male | 31 |
| XY-M-6 | XY | Male | 31.5 |

### Refining the PAR boundary

The boundary of the pseudoautosomal region (PAR), where the X and Y chromosomes still recombine, was previously estimated based upon genetic linkage maps s of the sex chromosomes [45,46] and patterns of molecular divergence between the X and Y chromosomes [19]. To determine which genes are within the PAR and which are in the X- and Y-unique regions, we compared expression of genes on the X and Y chromosomes between XX and XY fish. We summed normalized read counts across all tissues and samples for XX and XY samples respectively. We then calculated the log_2_ ratio of XY/XX expression for each gene (Figure 3.1). Both X and Y chromosome genes in the PAR have a log ratio centered around 0 as predicted for genes present in two copies in both XX and XY fish. Genes in the X-unique region have a log ratio centered around −1, consistent with a 2:1 dosage ratio for XX to XY fish. Genes in the Y-unique region have substantially elevated log ratios, supporting their absence from XX fish.

**Figure 3.1.**
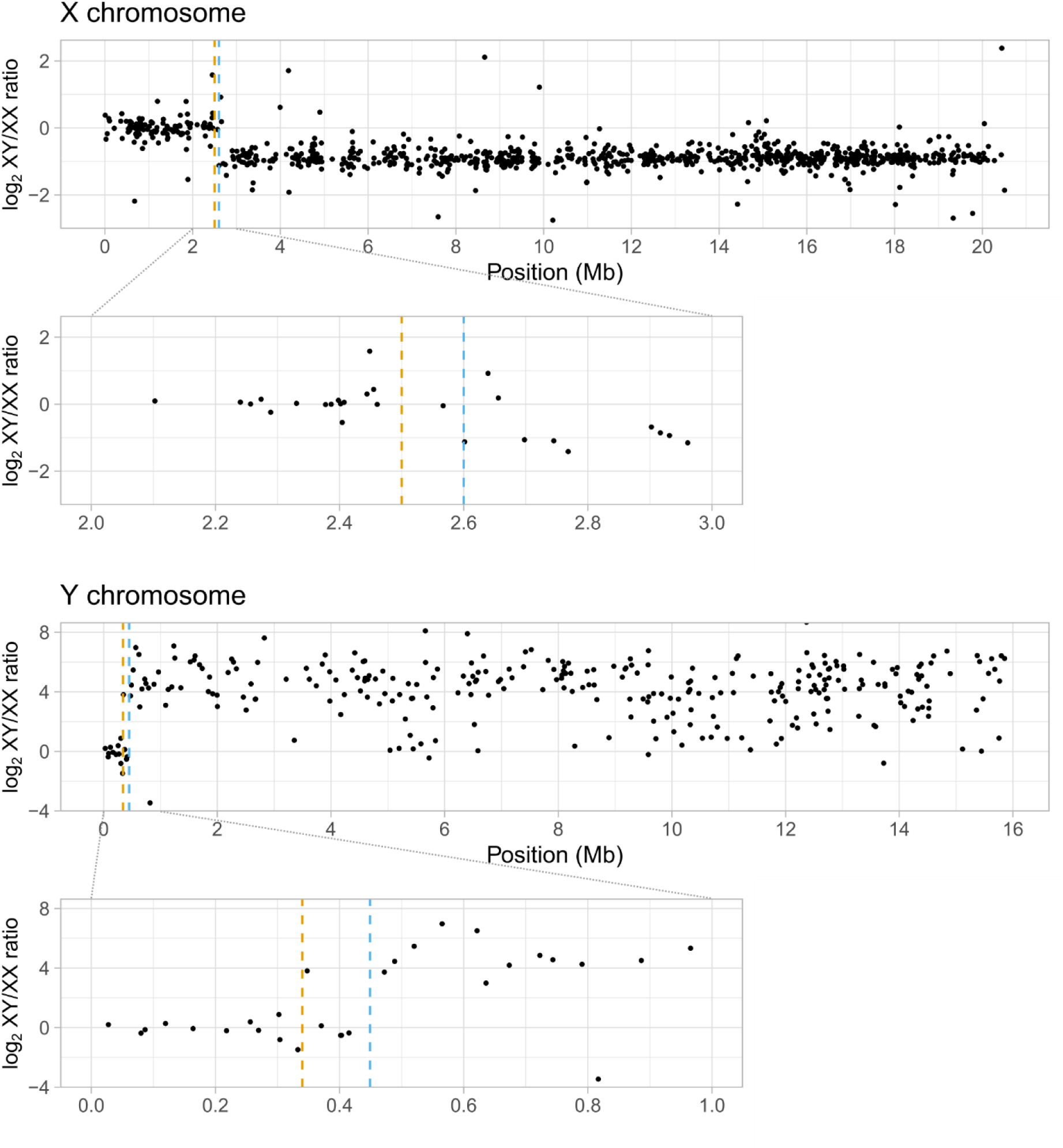
Expression of X and Y chromosome genes in XX and XY fish. Dot plots show log_2_ ratios of XY/XX normalized read counts across all tissues in males and females. Values near 0 indicate a 1:1 expression level between XX and XY genotypes which would be expected for genes within the PAR. Values of -1 indicate a 2:1 expression level which is expected for genes in the X unique region. Values greater than 1 indicate that the genes are within the Y unique region and are not present in XX genotypes. Previously defined PAR boundaries at 0.34 Mb on the Y chromosome and 2.5 Mb on the X chromosome are indicated by a vertical orange line. The PAR boundaries used in this study at 0.449 Mb on the Y chromosome and 2.601 Mb on the X chromosome are indicated in blue.

Expression ratios match previous estimates of the PAR boundary of 2.5 Mb on the X chromosome [18,19,45,46]. The PAR is not fully assembled on the threespine stickleback Y chromosome[17]. Our expression data indicates that the first 0.44 Mb of the Y chromosome is pseudoautosomal. We queried the longest transcripts of ten genes surrounding the putative Y boundary against all transcripts in the stickleback genome using BLAST (Table 3.2). The first six genes share high sequence identity with syntenic genes near the estimated X chromosome PAR boundary consistent with ongoing recombination between the X and Y chromosomes in the PAR. The sixth gene, LOC120812456, has a substantially shorter transcript length compared to its X chromosome match. This gene may have been truncated at an inversion breakpoint on the Y chromosome. The remaining four genes have poorer matches far from the PAR boundary, indicating that they are located within the non-recombining region of the Y chromosome. From these results, we set the PAR boundary at 0.449 Mb on the Y chromosome and 2.601 Mb on the X chromosome, placing 50 of the 692 annotated Y chromosome genes and 184 of the 1227 annotated X chromosome genes within the PAR. We masked the PAR on the Y chromosome and recalculated raw and normalized read counts for further analyses. Genes within the PAR are included with autosomal genes for the remaining results. X-linked, Y-linked, and sex-linked genes refers to genes within the nonrecombining region of the X, Y, or either sex chromosome, respectively.

**Table 3.2.** BLAST results of Y chromosome genes surrounding the proposed PAR boundary.

| Y Query |  |  |  | BLAST results |  |  |  |  |  |  |
| --- | --- | --- | --- | --- | --- | --- | --- | --- | --- | --- |
| Gene | Transcript length (bp) | Position | Gene type | Gene | Transcript length (bp) | Chromosome | Position | Gene type | Query coverage | Percent identity |
| LOC120812827 (rna-XM_040169038.1) | 1350 | 401732..403081 | protein coding | LOC120809745 (rna-XM_040163776.1) | 1366 | 19 | 2553601..2554966 | protein coding | 1.00 | 99.7% |
| LOC120812829 (rna-XM_040169039.1) | 588 | 403084..403671 | protein coding | LOC120809747 (rna-XM_040163777.1) | 610 | 19 | 2554969..2555578 | protein coding | 1.00 | 99.5% |
| LOC120812455 (rna-XM_040168425.1) | 1732 | 406481..411817 | protein coding | LOC120809573 (rna-XM_040163474.1) | 1831 | 19 | 2556895..2563599 | protein coding | 0.99 | 99.4% |
| LOC120812454 (rna-XM_040168422.1) | 3188 | 415112..448312 | protein coding | LOC120809049 (rna-XM_040162594.1) | 3219 | 19 | 2566894..2600202 | protein coding | 1.00 | 98.8% |
| LOC120812457 (rna-XR_005710621.1) | 1185 | 444593..447213 | lncRNA | LOC120809050 (rna-XR_005710068.1) | 1179 | 19 | 2596496..2599102 | lncRNA | 0.99 | 99.9% |
| LOC120812456 (rna-XR_005710620.1) | 1093 | 449610..452657 | lncRNA | nlrc5 (rna-XM_040162590.1) | 6665 | 19 | 2601499..2633055 | protein coding | 0.92 | 99.9% |
| LOC120812632 (rna-XR_005710686.1) | 1274 | 472268..488298 | lncRNA | LOC120812970 (rna-XR_005710787.1) | 598 | 2 | 1464441..1465308 | lncRNA | 0.46 | 94.9% |
| LOC120812631 (rna-XM_040168761.1) | 2564 | 488739..492589 | protein coding | cdkn1bb (rna-XM_040162614.1) | 2460 | 19 | 19575083..19578725 | protein coding | 0.51 | 91.2% |
| LOC120812429 (rna-XM_040168365.1) | 4065 | 520151..565054 | protein coding | snx1a (rna-XM_040163553.1) | 4313 | 19 | 15882890..15891931 | protein coding | 0.90 | 89.7% |
| LOC120812430 (rna-XR_005710617.1) | 1007 | 565219..567025 | lncRNA | LOC120808586 (rna-XR_005711822.1) | 2020 | 19 | 16730556..16738585 | lncRNA | 0.22 | 90.9% |

### Sex chromosomes have a larger effect on gene expression than gonadal sex in somatic tissues

We removed tRNAs and low expression genes (those with fewer than ten samples with at least five raw reads). This threshold retained 20749/26250 non-tRNA genes (19602/24531 autosomal, 854/1033 X-linked, and 293/686 Y-linked). We limited most analyses to autosomal genes, as dosage differences between the X and Y chromosomes cause the majority of sex-linked genes to be differentially expressed in all tissues between XX and XY genotypes, inflating the apparent effect of sex chromosomes on genome-wide expression differences.

We analyzed the independent effects of gonadal sex and sex chromosome complement on gene expression to compare the relative contribution of each factor to sex differences in gene expression. We found no clear separation of somatic tissues by gonadal sex or sex chromosomes through principal component analysis (PCA) of autosomal gene expression (Figure 3.2), indicating an overall similarity in gene expression within each tissue irrespective of sex. When comparing the number of autosomal differentially expressed genes (DEGs) associated with each factor, all somatic tissues had more DEGs in the sex chromosome comparison (XX/XY) than the gonadal sex comparison (Female/Male) (Table 3.3). We examined the magnitude of the effect of both factors on gene expression by comparing the absolute value of the shrunken log_2_ fold change (|LFC|) for all autosomal genes. The shrinkage transformation reduces the LFC for high dispersion genes, making more conservative estimations of effect size that is less impacted by noisy and low count genes [47]. In all three tissues, XX/XY |LFC| was significantly higher than Female/Male |LFC| (Figure 3.3, Table 3.4). Taken together, the number of DEGs and average effect sizes in brain, gill, and liver all indicate that sex chromosome complement has a larger effect on overall gene expression differences than gonadal sex in somatic tissues. The majority of DEGs did not overlap between the Female/Male and XX/XY comparisons in somatic tissues, indicating that genes tend to be downstream of either sex-linked genes or regulation of gonadal hormones, but not both factors.

**Figure 3.2.**
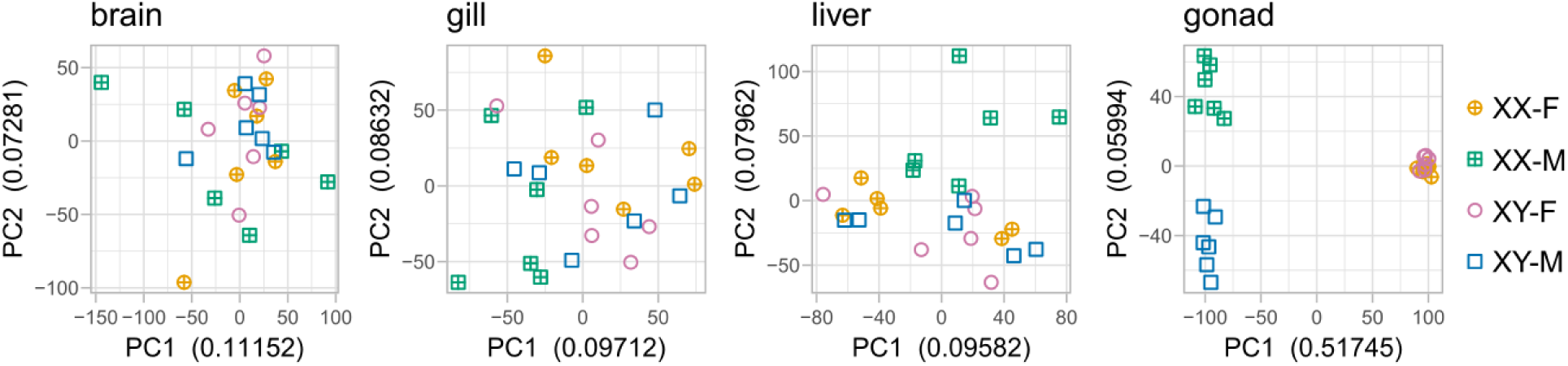
PCA plots of normalized read counts for autosomal genes within each tissue. Principal Components 1 and 2 with the Proportion of Variance Explained are shown on the X and Y axis respectively. In the gonad, samples separate by gonadal sex on PC1 and by sex chromosome on PC2. In somatic tissues, samples do not show any clear separation by genotype.

**Figure 3.3.**
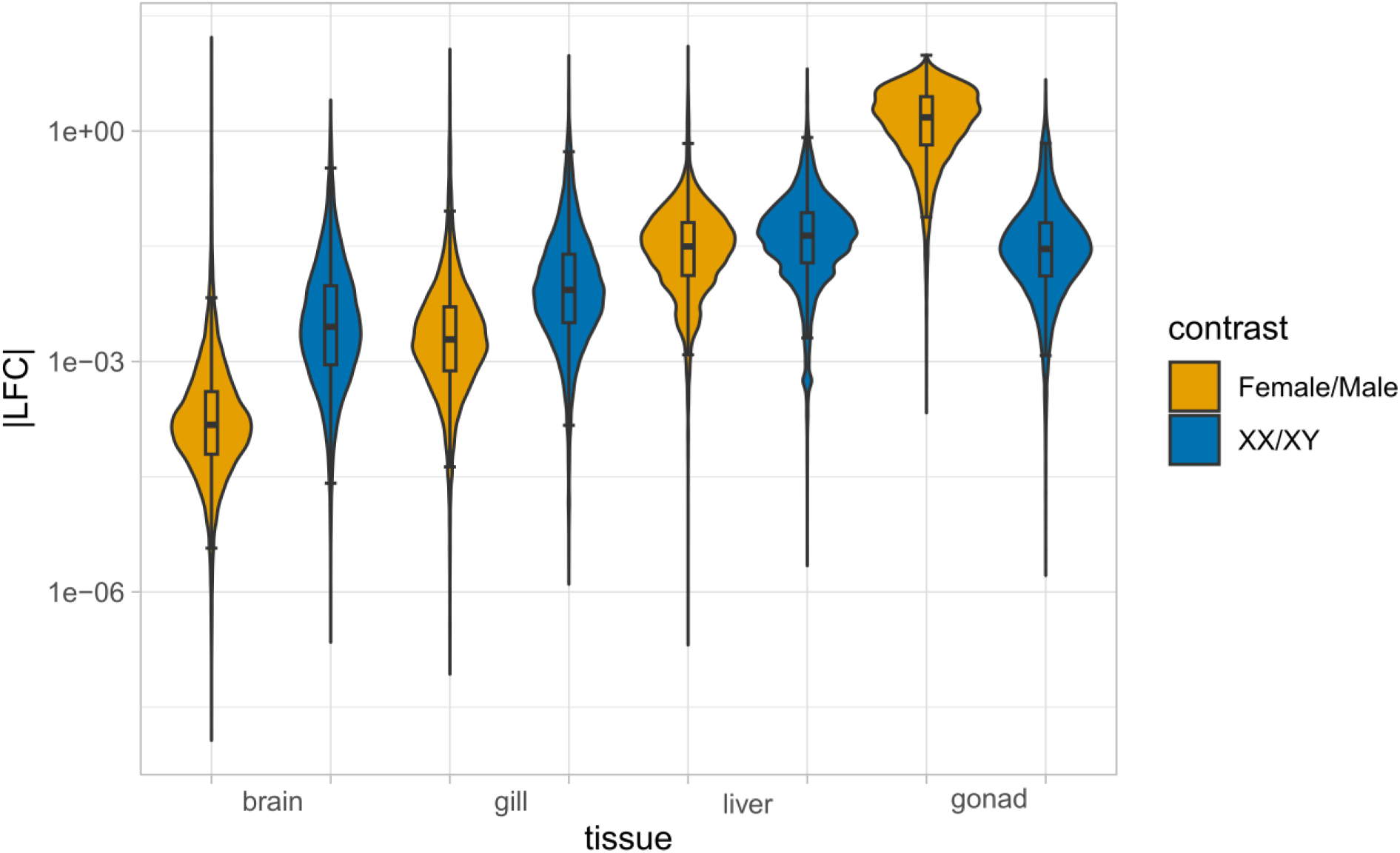
|LFC| distributions for autosomal genes in each tissue. Shrunken |LFC|, which represents the magnitude of the effect of gonadal sex or sex chromosomes, of all autosomal genes are shown for Female/Male contrast in orange and XX/XY contrast in blue.

**Table 3.3.** Total number of DEGs within each tissue. The portion of DEGs for each tissue that are within each group is shown in parentheses.

| Tissue | Total DEGs | Female/Male DEGs | XX/XY DEGs | Both Female/Male and XX/XY DEGs |
| --- | --- | --- | --- | --- |
| Brain | 115 | 11 (0.0957) | 107 (0.930) | 3 (0.0261) |
| Gill | 315 | 77 (0.244) | 269 (0.854) | 31 (0.0984) |
| Liver | 740 | 279 (0.377) | 516 (0.697) | 55 (0.0743) |
| Gonad | 16190 | 16108 (0.995) | 697 (0.0431) | 615 (0.0380) |

The brain, which had the fewest number of autosomal DEGs, showed the greatest difference in DEG count with tenfold more XX/XY DEGs than Female/Male DEGs. This ratio decreased across tissues as the number of total DEGs increased, with XX/XY DEGs 3.5-fold higher in gill and 1.8-fold higher in liver. When comparing across tissues, Female/Male DEGs showed more variation in number (25-fold difference between brain and liver) compared to XX/XY DEGs (5-fold difference between brain and liver). This indicates that response to gonadal hormones is more variable across tissues, whereas sex chromosome driven differences are more consistent.

### Gonadal sex has a larger effect on gene expression than sex chromosomes in the gonad

The gonad showed contrasting patterns in the effect of sex chromosomes and gonadal sex compared to somatic tissues. In the gonad, PCA of autosomal expression shows clear separation by sex on PC1, with separation by sex chromosome present in PC2 (Figure 3.2). There were substantially more Female/Male DEGs than XX/XY DEGs (Table 3.3), with 85% of the 19062 autosomal genes remaining after filtering differentially expressed between Males and Females. |LFC| was dramatically higher for Female/Male DEGs than XX/XY DEGs (Figure 3.3, Table 3.4). These results are not surprising, given the extensive differences in cellular composition and function of the ovary and testis, relative to the similarities in structure of somatic tissues between males and females. While sex chromosomes have a lesser effect than sex in the gonad, the gonads have more XX/XY DEGs than any somatic tissue. XX/XY |LFC| in the gonad are higher than in the brain or gill, but lower than |LFC| in the liver. This again shows a relatively consistent effect of sex chromosomes on overall gene expression across tissues, even in the presence of dramatic tissue composition and expression differences that are observed in the gonad.

**Table 3.4.** Mean and median LFC values for all autosomal genes. P-values are shown for two-sided binomial sign test between the Female/Male and XX/XY LFC. Significant values are bolded (p = 0.05).

| Tissue | Contrast | Mean LFC | Median LFC | p-value |
| --- | --- | --- | --- | --- |
| Brain | Female vs Male | 0.0029 | 0.0001 | < **2.2e-16** |
|  | XX vs XY | 0.0177 | 0.0028 |  |
| Gill | Female vs Male | 0.0128 | 0.0019 | < **2.2e-16** |
|  | XX vs XY | 0.0397 | 0.0085 |  |
| Liver | Female vs Male | 0.0622 | 0.0315 | < **2.2e-16** |
|  | XX vs XY | 0.0835 | 0.0435 |  |
| Gonad | Female vs Male | 1.8925 | 1.5086 | < **2.2e-16** |
|  | XX vs XY | 0.0745 | 0.0292 |  |

### The X and Y chromosome are enriched for Female/Male DEGs

By comparing all four genotypes, we can control for dosage differences of X- and Y-linked genes and evaluate the effect of gonadal sex on their expression. In all somatic tissues, a higher proportion of X-linked and Y-linked genes remaining after filtering are Female/Male DEGs compared to autosomal genes (Figure 3.4, Table 3.5). In somatic tissues, both X- and Y-linked DEGs were Female biased more frequently than autosomal DEGs, though this difference is not significant. In the gonad, the X chromosome was not enriched for Female/Male DEGs, and sex bias was similar to autosomal DEGs. The Y chromosome had significantly fewer Female/Male DEGs than the autosomes, but these DEGs were male biased significantly more often than autosomal DEGs. These findings are consistent with predictions that sex chromosomes accumulate genes with sex specific functions [1,4].

**Figure 3.4.**
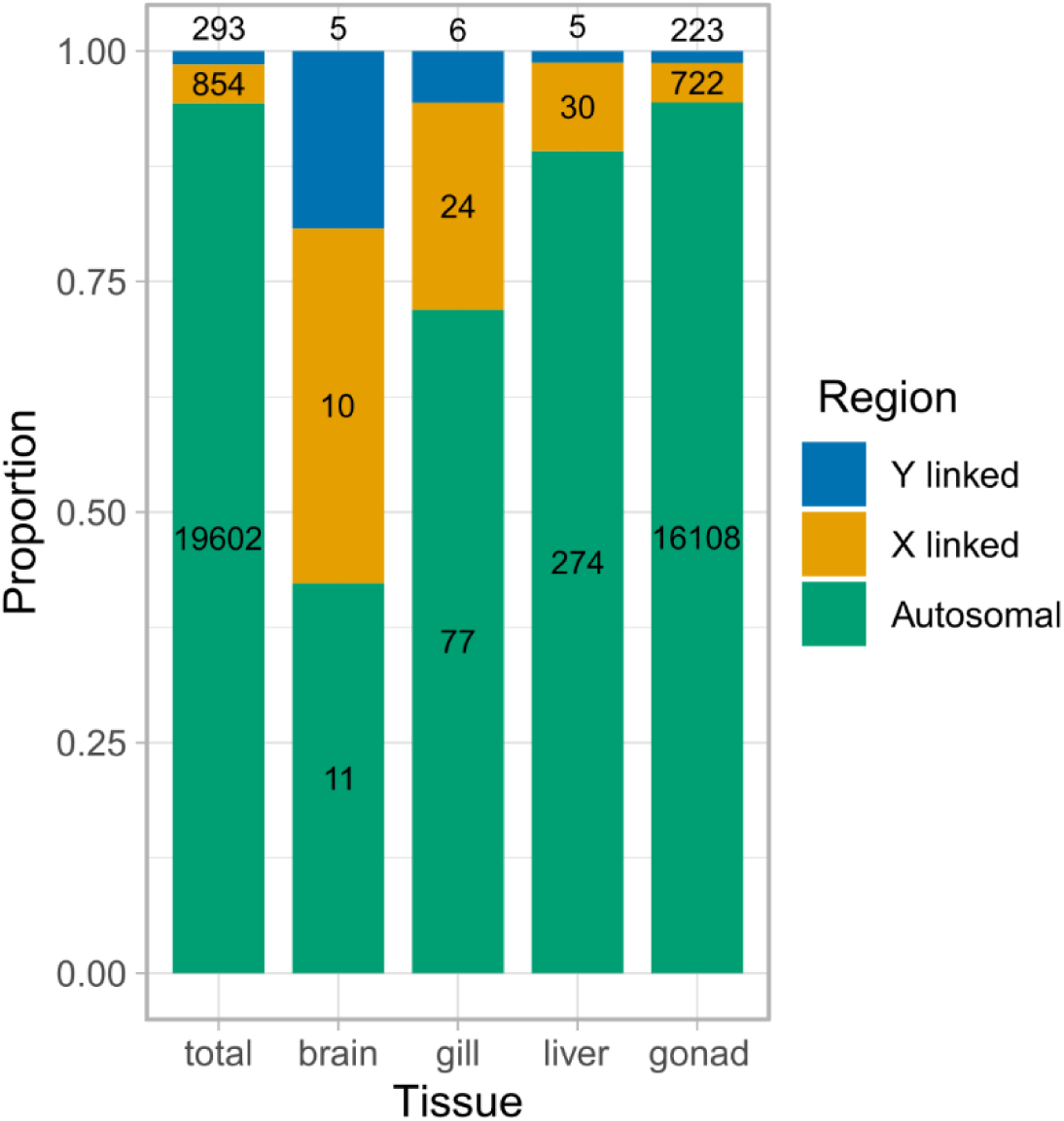
Proportion of Female/Male DEGs in different regions of the genome. Total column shows the number of genes remaining in each region after filtering out low count genes. Other columns show the number of DEGs in each region. The number of genes in each category is shown within or above the bars.

**Table 3.5.** Female/Male DEGS are enriched on X and Y. The DEGs column shows the number of DEGs from each genomic region and the proportion genes that are DEGs within that region (Total number of genes remaining after filtering: 19602 autosomal genes, 854 X-linked genes, and 293 Y-linked genes total). Female and Male biased columns show the number and proportion of DEGs with the respective bias. P-values show Fisher’s exact test compared to the proportion of autosomal genes that are differentially expressed or the proportion of autosomal DEGs that are Female or Male biased.

| Tissue | Region | DEGs | p-value | Female biased | Male biased | p-value |
| --- | --- | --- | --- | --- | --- | --- |
| Brain | Autosomal | 11<br>(0.000578) | - | 4<br>(0.364) | 7<br>(0.636) | - |
|  | X-linked | 10<br>(0.0117) | **4.58e-09** | 6<br>(0.600) | 4<br>(0.400) | 0.395 |
|  | Y-linked | 5<br>(0.0171) | **2.93e-06** | 3<br>(0.600) | 2<br>(0.400) | 0.596 |
| Gill | Autosomal | 77<br>(0.00404) | - | 31<br>(0.403) | 46<br>(0.597) | - |
|  | X-linked | 24<br>(0.0281) | **4.93e-12** | 15<br>(0.625) | 9<br>(0.375) | 0.0645 |
|  | Y-linked | 6<br>(0.0205) | **1.63e-03** | 4<br>(0.667) | 2<br>(0.333) | 0.2353 |
| Gonad | Autosomal | 16108<br>(0.845) | - | 5572<br>(0.346) | 10536<br>(0.654) | - |
|  | X-linked | 722<br>(0.845) | 1.00 | 257<br>(0.356) | 465<br>(0.644) | 0.576 |
|  | Y-linked | 223<br>(0.761) | **2.39e-04** | 57<br>(0.256) | 166<br>(0.744) | **0.00449** |
| Liver | Autosomal | 274<br>(0.0144) | - | 141<br>(0.515) | 133<br>(0.485) | - |
|  | X-linked | 30<br>(0.0351) | **4.56e-05** | 16<br>(0.533) | 14<br>(0.467) | 1.00 |
|  | Y-linked | 5<br>(0.0171) | 1.00 | 4<br>(0.8) | 1<br>(0.200) | 0.372 |

### Most DEGs are tissue specific

To further evaluate the similarity of the effect of sex factors across tissues, we compared which genes were differentially expressed within each tissue (Figure 3.5). XX/XY autosomal DEGs are largely nonoverlapping, with between 78.9% and 92.7% of DEGs of a given tissue being tissue specific. Most X-linked and Y-linked XX and XY DEGs are DE in all tissues, which is expected given the dosage differences between XX and XY genotypes and the lack of a chromosome level dosage compensation mechanism in threespine stickleback [19,20]. Female/Male autosomal DEGs in somatic tissues overlap substantially with the gonad; however, this is likely not meaningful, as the vast majority of all genes are Female/Male DEGs within the gonad. Excluding the gonad, 90.9% to 99.6% of Female/Male autosomal DEGs within each somatic tissue are tissue specific. Very few Y-linked genes are differentially expressed between Females and Males in somatic tissues, but most Female/Male X-linked DEGs are tissue specific across somatic tissues, similar to the pattern seen with autosomal DEGs. In total, the specific genes which are impacted by both gonadal sex and sex chromosome complement are highly tissue dependent.

**Figure 3.5.**
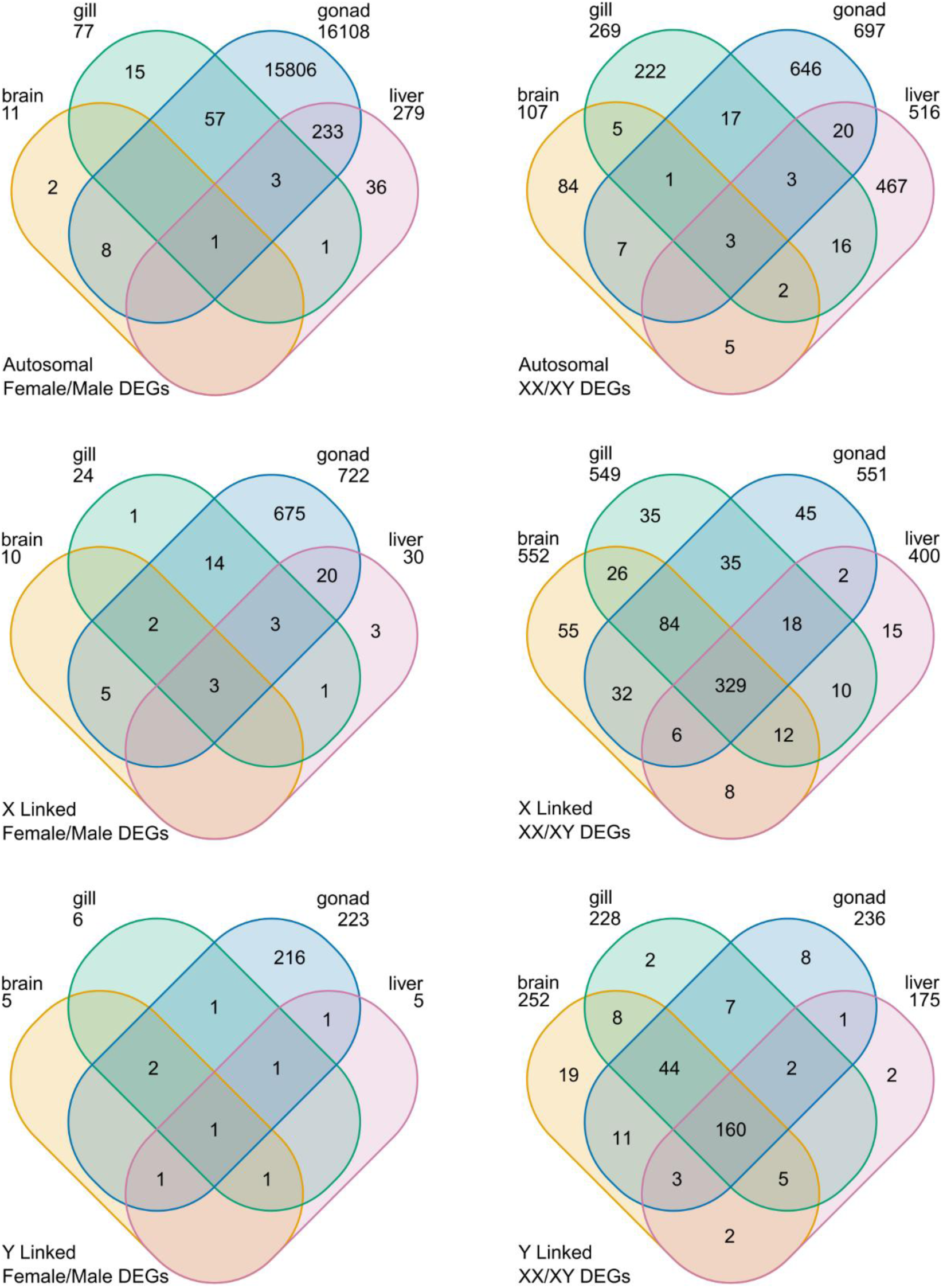
Overlap of DEGs between different tissues. The total number of DEGs are shown below each tissue.

### The *amhy* transgene shows elevated expression relative to the endogenous locus

The *amhy* transgene in XX-M fish utilizes native putative regulatory elements from the Y chromosome; however, expression patterns in transgenes are often influenced by where the transgene integrates into the genome [48,49]. To determine whether the *amhy* transgene may have insertion specific effects on expression, we compared expression of *amhy* from the transgene to the endogenous locus on the Y chromosome. We observed elevated expression of *amhy* in transgenic XX-M relative to XY-M in gonad, brain, and gill tissue (Figure 3.6). Expression in the liver was minimal in both genotypes. Since *amhy* and its autosomal paralog, *amh*, both likely utilize the same dedicated receptor, *amhrII*, we also examined expression of these genes to contextualize potential effects of ectopic *amhy* expression. Of note, the poly(A) based mRNA sequencing we have used is not biased by transcript length [50,51], allowing direct comparison of normalized read counts between different genes. Expression of *amhrII* was negligible in all somatic tissues when compared to the gonad, which would likely limit the effects of ectopic *amhy* expression in somatic tissues. Expression of *amh* was also low in somatic tissues of both genotypes which is consistent with *amh* expression patterns in other taxa. In the gonad, *amh* expression in the gonad exceeded *amhy* in either genotype, which may dampen differences in *amh* signaling caused by excess *amhy* in XX-M. However, *amh* expression is elevated in XX-M compared to XY-M, which could indicate subtle developmental differences between these two genotypes or an effect of elevated *amhy* expression in the gonad.

**Figure 3.6.**
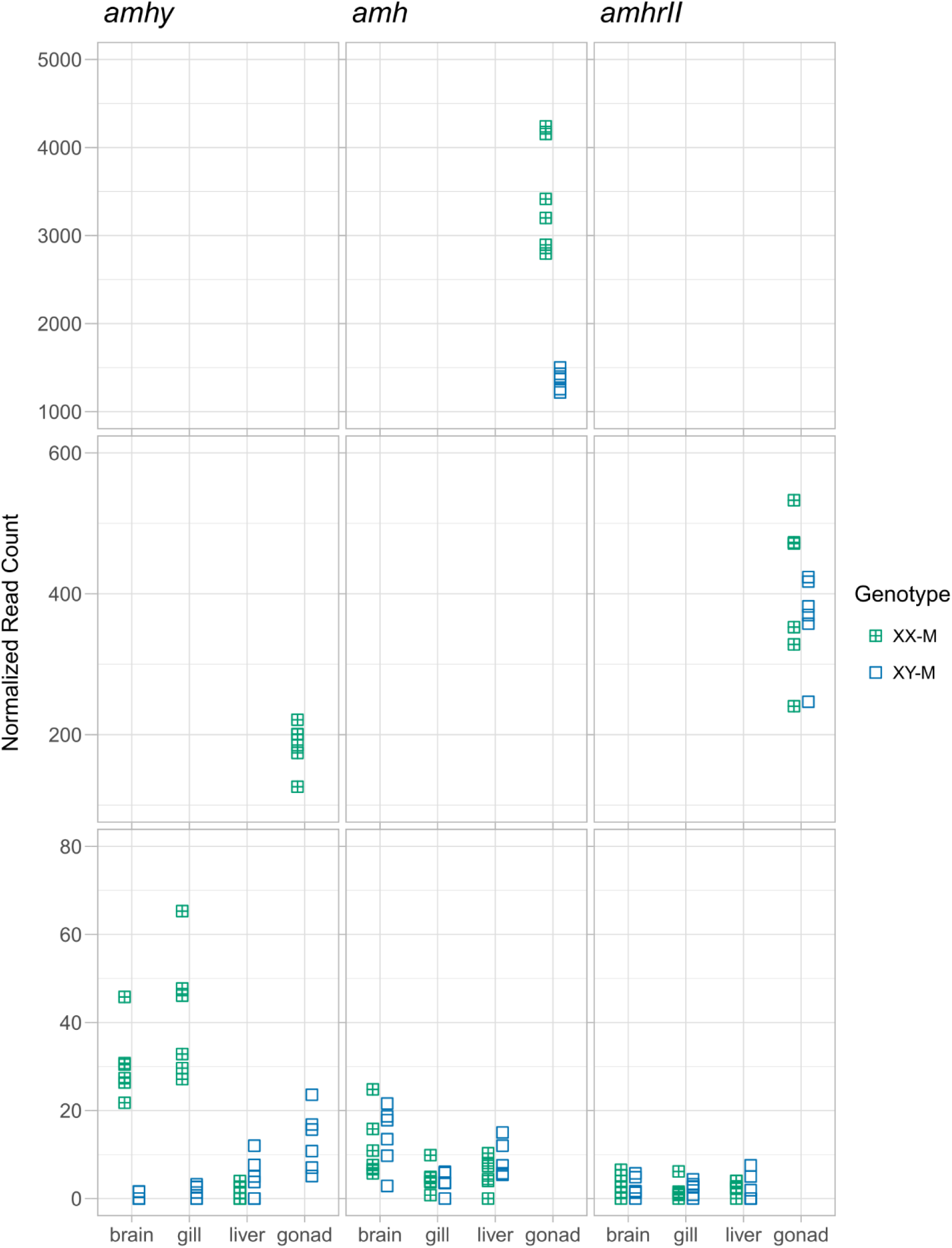
Expression of *amhy*, *amh*, and *amhrII* in wildtype and sex reversed males. Normalized read counts of the sex determination gene, *amhy*, its autosomal paralog, *amh*, and the dedicated receptor, *amhrII*, in sex reversed XX-M and wildtype XY-M are shown for each tissue. Note the changes in Y axis scales at 80 and 600 units.

## Discussion

### Gonadal sex and sex chromosomes have distinct effects on gene expression

We find that sex chromosomes have a substantial impact on sexually dimorphic gene expression in the threespine stickleback. The effect of sex chromosomes is most striking in somatic tissues, where sex chromosomes affect the expression of more genes and have a larger effect size than gonadal sex. While gonadal sex clearly has a larger effect than sex chromosomes in the gonad, the effects of sex chromosomes on gene expression in the gonad is not insignificant. The gonads have more XX/XY DEGs than any somatic tissue, and the gonad is the only tissue to show any separation by sex chromosome in PCA. This is consistent with the expectation that sex chromosomes will accumulate genes with gonad specific function [1,4]. The overwhelming effect of gonadal sex on gene expression in the gonad is not surprising, given the vast physiological differences between the ovary and testis which would be captured only in the Female/Male contrast of this tissue. Likewise, ovaries and testes are expected to show far more expression differences than somatic tissues, where differences between females and males would be primarily due to differences in sex hormones produced by the gonad.

When compared to the Four Core Genotypes mouse model, sex chromosome complement appears to have a larger contribution to gene expression differences in stickleback than in mouse. In mouse, gonadal sex is shown to have a larger effect on gene expression in the liver [52] and hippocampus [53]. One might expect the more diverged mammalian sex chromosomes to give rise to more expression differences than the less diverged sex chromosomes in stickleback. However, the complete dosage compensation present in mammals offsets the difference in gene content between XX and XY genotypes very effectively, and few X-linked genes show differential expression by sex chromosome complement in mouse [52,53]. In contrast, threespine stickleback lack chromosome-level dosage compensation [19,20], and the majority of X chromosome genes are differentially expressed between XX and XY fish. While an XY mouse is hemizygous for far more genes than an XY stickleback, stickleback have a larger difference in effective dosage of X chromosome genes. Thus, it is reasonable that sex chromosome complement would have a larger effect on expression of autosomal genes in the threespine stickleback than in the mouse.

Reproductive maturation may also contribute to the larger role of the sex chromosomes we have observed. A time course analysis of expression in the mouse heart shows that sex chromosomes affect a similar number of genes in 16.5 dpc embryos, neonates, and adults [54]. Gonadal sex has a negligible effect at 16.5 dpc and in neonates, but greatly exceeds the effect of sex chromosomes in adults [54]. Relative to the juvenile fish we analyzed in this study, the effect of gonadal sex on gene expression would likely be higher in reproductively mature fish when circulating sex hormones are at their peak, which occurs around one year of age in wild populations [55,56]. It would also be informative to measure hormone levels in all four genotypes throughout development to assess how closely sex reversed fish recapitulate hormone levels of their wildtype counterparts.

### Sex biases of sex chromosomes

Typically, expression comparisons of X- and Y-linked genes between males and females are confounded by dosage differences between the XX and XY genotypes and can be challenging to interpret. However, a Four Core Genotypes system provides the unique opportunity to control for these dosage differences and measure the effect of gonadal sex on these genes. We saw that both X- and Y-linked genes are significantly more likely to be affected by gonadal sex than autosomal genes. In somatic tissues, this suggests that sex-linked genes have evolved regulatory elements responsive to, or be downstream of genes responsive to, sex hormones produced by the gonad. Alternatively, hormone responsive genes may have been enriched on the ancestral autosome or were more likely to be retained on the sex chromosomes once recombination was suppressed. For X-linked genes, these elements could compensate for dosage differences between the XX and XY genotypes, leading to male biased expression patterns after normalizing for sex chromosome complement. Alternatively, if dosage differences between males and females were beneficial, hormone responsive elements could act in the same direction as dosage differences, amplifying an already female biased expression. While the total number of sex-linked DEGs is low, limiting statistical power, we see similar numbers of female and male biased DEGs on the X chromosome. This indicates that both compensating and amplifying hormone responses exist and operate differently depending on the target gene. Y-linked genes are only present in the male background, so one might not expect them to utilize differences in hormonal cues to regulate their expression. However, androgen and estrogen signaling could help to synchronize expression of these genes with sexual maturation, facilitating some sex specific function.

Interpreting these differences is more challenging in the gonad due to the overwhelming differences in structure, function, and autosomal gene expression between these two tissues. X-linked genes can become femininized in function, as the X chromosome is in females more often than in males [57–59]. Conversely, X-linkage can favor male specific functions, as recessive X alleles are always exposed in the hemizygous XY genotype, enhancing the strength of selection [60–63]. Both X and Y linked genes are often important to reproductive development and fertility [64–66]. Sex reversed XY-F stickleback have a noticeable reduction in fertility, but we do not yet know if this is due primarily to reduced dosage of key X-linked factors or interference from Y-linked genes not typically present in females. Males do not show obvious fertility defects, so the threespine Y chromosome clearly does not carry any factors essential for spermatogenesis or male fertility as is seen in drosophila [64] and mammals [65]. However, sex-linked loci may have more subtle effects on these traits which could be further canalized as the X and Y continue to diverge. Further research into the effects of these sex-linked genes will provide valuable insight into how sex chromosomes evolve sex specific functions in reproductive development.

### Limitations

We observed elevated expression of *amhy* in transgenic XX-M compared to wildtype XY-M in the gonads and in somatic tissues. While the *amhy* transgene utilizes endogenous putative elements on the Y chromosome, this finding is not surprising and is likely influenced by position effects, where the location of a transgene within the genome influences its expression [48,49]. The impact of excess *amhy* is likely to be limited in somatic tissues due to the low levels of its dedicated receptor, *amhrII*. There is evidence in zebrafish, which lack *amhrII,* that *amh* uses the type II TGF-β receptor *bmpr2a* [67,68], so we should not rule out interactions between *amhy* and noncanonical receptors. In the gonad, where *amhrII* is highly expressed, expression of *amhy* from the transgene is much lower than expression of the autosomal paralog *amh,* which would likely buffer the effect of elevated *amhy* expression from the transgene.

Another limitation in this study is the cross scheme. As this experiment utilizes the first generation of F1 XX-M fish, these fish were made in a separate cross from the other genotypes. Differences in genetic background from our outbred population and minor differences in rearing conditions across clutches could confound comparisons. We are in the process of crossing the *amhy* transgene into a line of fish with the loss of function *amhy* allele used in this study. This will allow the sex determining *amhy* transgene to segregate independently of the sex chromosomes. Using an XY^amhy−^ *Tg(amhy)* male, a single cross with a wildtype XX female would be able to generate all four genotypes, eliminating this limitation in future studies.

### Future directions

We have shown that sex chromosome complement has a significant contribution to sexually dimorphic gene expression in the threespine stickleback. Whether these expression differences give rise to phenotypic differences that could impact fitness remains to be seen. Threespine stickleback have a wide range of well documented mating behaviors and also exhibit male only parental care. Expression differences in the brain caused by sex chromosome complement could play an important role in these essential behaviors. There are also many anatomical differences between males and females, including bony armor features which protect from predation [22,23] and size and shape of the head and body [22–25]. The sex specific fitness consequences of these traits are less clear, but as these characteristics would be established long before reproductive maturity, intrinsic genetic differences that are present from fertilization may be a more reliable avenues to encode these traits than hormonal control. Alternatively, the expression differences we have observed could be neutral or even deleterious results of Y chromosome degeneration which have yet to be solved by a chromosome level dosage level mechanism. In general, genes that are predicted to be dosage sensitive have been conserved on the Y [19], but as more of these genes degenerate, selective pressure for dosage compensation both at the gene and chromosome level could accrue. Further work with the Four Core Genotypes stickleback will be invaluable for uncovering the role in sex chromosomes in sex differences and how these differences, in turn, shape sex chromosome evolution.

## Methods

### Animal Husbandry

All experiments used lab-derived progeny from wild-caught threespine stickleback fish from Port Gardner Bay (Washington, USA). All procedures were approved by the University of Georgia Animal Care and Use Committee (protocols A2021 07-031-Y3 and A2024 08-009-Y1). We collected fish from Washington State over multiple years using Scientific Collection Permit numbers 21-122, 22-150, 23-147, and 24-143. All experiments were performed using fish crossed and reared in laboratory conditions. Fish were maintained in 3.5 ppt Instant Ocean Sea Salt (Spectrum Brands, Blacksburg, VA, USA) in reverse osmosis water at 18 °C and pH 8.0 and a summer photoperiod (16L:8D).

### Tissue collection

We generated mutant and wildtype fish from two sets of crosses. To obtain sex reversed XY females (XY-F) and wildtype XX females (XX-F) and XY males (XY-M), we crossed one wildtype XY-M with an XY-F carrying a previously described +9,+5 *amhy*-KO allele [44]. This cross yields wildtype XX-F and XX-M fish which receive the maternal X chromosome as well as sex reversed XY-F fish and embryonic lethal YY fish which inherit the maternal Y chromosome. We generated sex reversed XX males (XX-M) from one F0 XX-M with germline integrations of the -5*amhy*2.5,Xla.Ef1a:EGFP transgene heritable to 7% of offspring [44]. This transgene contains *amhy* under control of its putative native regulatory elements and a ubiquitously expressed EGFP reporter [44]. We crossed the XX-M with three wildtype XY-F. At five days post fertilization, we identified transgenic F1 fish by expression of EGFP and discarded negative embryos.

We euthanized fish in 0.05% MS-222 buffered to neutral. We dissected Tissues and preserved them in RNAlater Stabilization Solution (Invitrogen AM7021) following manufacturer guidelines and stored at -20 °C until analysis. We extracted RNA using TRIzol Reagent (Invitrogen 15596026) and the Direct-zol RNA Microprep Kit (Zymo R2062). We extracted DNA from fin clips using a HotSHOT isolation protocol [69]. We performed all PCR reactions on an Analytik Jena Biometra Tone in a total volume of 20 µl using 0.8 μL of genomic DNA, 0.2 μM each primer, 0.2 mM each dNTPs, 0.5 units DreamTaq DNA polymerase (Thermo Scientific EP0714), and 2 µl 10x DreamTaq Green Buffer with 20 mM MgCl_2_. All PCR reactions used the following cycling conditions: 1 cycle of 95°C for 2 min; 35 cycles of 95°C for 30 s, 60°C for 30 s, and 72°C for 30 s; 1 cycle of 72°C for 5 minutes. We genotyped fish for sex chromosome using the sex-linked marker *idh* [18] using primers 5’-CCACTTGCAGTTTGTCTGAAGG-3’ and 5’-TGACAGTCCTACTCAAGGCA-3’. We identified the *amhy* transgene and *amhy* KO allele as previously described [44], using primers for *EGFP* (5’-ATCATGGCCGACAAGCAGAA-3’ and 5’-AACTCCAGCAGGACCATGTG-3’) and *amhy* (5’-CAGCCGGAGTCAGTTTGGAA-3’ and 5’-TCATGCATCGTCGACTGGAG-3’).

### RNAseq

Libraries were prepared and sequenced by Admera Health (South Plainfield, NJ USA). Libraries were prepared using the Alithea Genomics Mercurius BRB-seq kit. 1.16B 150 bp paired end reads were sequenced on an Illumina NovaSeq X plus. We filtered R2 reads with Fastp v0.23.4 [70] using default quality thresholds with the addition of the following parameters: *--trim_poly_g --trim_poly_x --cut_right --length_required 25*. We did not filter R1 reads, as they are used only for barcode identification and are not used for genome alignment. 880M reads were left after quality filtering, of which 854M had valid barcodes. We aligned reads to the threespine stickleback reference genome (GAculeatus_UGA_version5, NCBI RefSeq assembly GCF_016920845.1) and generated counts tables using the alignReads and GeneCounts function in STAR v2.7.11b [71] following guidelines from the Alithea BRB-seq manual, resulting in 691M uniquely mapped reads. We performed all remaining analyses in R (v. 4.5.1) and RStudio (v. 2025.09.2+418). We removed all tRNA genes and genes without at least ten samples with at least five raw reads. We used DESeq2 (v1.48.1) [72] to generate normalized read counts and perform differential expression analysis. We identified DEGs using the standard workflow with the following design: *∼ tissue + sex + chromosome + tissue:sex + tissue:chromosome*. We used the contrast function to calculate log2 fold changes (LFC) and adjusted p values for differential expression between Females and Males and between XX and XY chromosomes in each tissue. LFC shrinkage was performed using the package ashr (v2.2-63) [47].

### BLAST

We retrieved mRNA sequences for all genes in the threespine assembly GAculeatus_UGA_version5 (GCF_016920845.1) using SAMtools v1.21 [73] and gffread v0.12.7 [74]. We queried Y transcripts listed in Table 3.X against other transcripts using BLAST+ v2.16.0 [75] with the following parameters: *-task dc-megablast -max_target_seqs 50 - max_hsps 5 -evalue 1e-3 -qcov_hsp_perc 10 -perc_identity 20*. We removed self-hits to any isoform of the query gene and selected the match with the highest bitscore for each query.

